# Retinal ganglion cell endowment is correlated with optic tract fiber cross section, not density

**DOI:** 10.1101/2022.03.30.486288

**Authors:** Huseyin O Taskin, Yuchuan Qiao, Valerie J Sydnor, Matthew Cieslak, Edda B Haggerty, Theodore D Satterthwaite, Jessica IW Morgan, Yonggang Shi, Geoffrey K Aguirre

## Abstract

There is substantial variation between healthy individuals in the number of retinal ganglion cells (RGC) in the eye, with commensurate variation in the number of axons in the optic tracts. Fixel-based analysis of diffusion MR produces estimates of fiber density (FD) and cross section (FC). Using these fixel measurements along with retinal imaging, we asked if individual differences in RGC tissue volume are correlated with individual differences in FD and FC measurements obtained from the optic tracts, and subsequent structures along the cortical visual pathway. We find that RGC endowment is correlated with optic tract FC, but not with FD. RGC volume had a decreasing relationship with measurements from subsequent regions of the visual system (LGN volume, optic radiation FC/FD, and V1 surface area). However, we also found that the variations in each visual area were correlated with the variations in its immediately adjacent visual structure. We only observed these serial correlations when FC is used as the measure of interest for the optic tract and radiations, but no significant relationship was found when FD represented these white matter structures. From these results, we conclude that the variations in RGC endowment, LGN volume, and V1 surface area are better predicted by the overall cross section of the optic tract and optic radiations as compare to the intra-axonal restricted signal component of these white matter pathways. Additionally, the presence of significant correlations between adjacent, but not distant, anatomical structures suggests that there are multiple, local sources of anatomical variation along the visual pathway.

## Introduction

The number of retinal ganglion cells (RGCs) in the eye varies two-to-threefold between healthy individuals (Curcio & Allen, 1990). As the RGCs give rise to the optic nerve and then optic tract, this variation is reflected directly in the number of axons within those nerves. Signals from the RGCs are expanded in pools of neurons within the lateral geniculate nucleus and then visual cortex, with a roughly 100x increase in the number of neurons in area V1 as compared to the retinal ganglion cell layer. There is also substantial individual variation in the size of these post-retinal visual pathway structures (Kupfer et al., 1967; Andrews et al., 1997; Aguirre et al., 2016). However, it is unknown how individual variation in RGC endowment is related to individual variation in the size of subsequent elements of the visual pathway.

In addition to direct volumetric measurement using structural magnetic resonance imaging (MRI), the white matter connections of the visual pathway have been studied using diffusion MR. A promising diffusion measure for the study of anatomical variation is fixel-based analysis (Raffelt et al., 2017). The fixel approach uses fiber orientation distribution maps (FOD) to extract separate estimates of fiber density (FD) and fiber cross section (FC). FD is ostensibly proportional to the intra-axonal volume of a fiber bundle (Raffelt et al., 2012), while FC is related to the cross-sectional area of the bundle. In patients with glaucoma, which results in loss of retinal ganglion cells, a decrease in FD and FC has been observed in the optic tract (Haykal et al., 2019). This finding in a patient population suggests that individual variation in retinal ganglion cell number in healthy subjects might be reflected in fixel measures of the post-retinal white matter pathway.

In recent work we have measured individual differences in the volume of the ganglion cell layer of the retina in forty-two people (Chen et al., 2020). These measures were adjusted for the confounding effect of differences in the axial length of the eye, yielding an estimate of RGC endowment for each subject. In these subjects we also obtained structural and diffusion MRI measures. Here we examine how individual variation in the RGCs is related to fixel-based measures of the optic tract and optic radiation. Secondarily, we consider how variation in anatomical measures along the visual pathway are related to one another, and to variation at the front-end of the visual system.

## Methods

### Subjects

42 normally sighted subjects from the University of Pennsylvania and surrounding Philadelphia community were studied under a pre-registered project (https://osf.io/ervrh). Participants were at least 18 years of age, had no history of ophthalmologic disease, and had corrected visual acuity of 20/40 or better. Fourteen of the subjects were women, and the mean age of all participants was 28. All participants provided written consent and the protocol was approved by the University of Pennsylvania Institutional Review Board. The retinal measurements used here are described fully in a prior publication (Chen et al., 2020).

### MRI acquisition

Diffusion, Blood Oxygen Level Dependent (BOLD), and anatomical images were acquired on a Siemens 3T Prisma scanner with a 64-channel head coil at the University of Pennsylvania using the HCP LifeSpan protocol (VD13D). The diffusion images were obtained with a multi-band Echo Planar Imaging (EPI) sequence, 1.5mm isotropic voxels, b= [0,1500,3000], number of gradient directions=98, acceleration factor=4, monopolar scheme, TR=3230ms, and TE=89.20ms. For the T1w images, the tfl3d1 sequence was used with 0.8mm isotropic voxels, TR=2400ms, TE=2.2ms, and flip angle=8°. For the T2w images the SPC sequence was used with 0.8 mm isotropic voxels, TR=3200ms, TE=563ms, flip angle=120°. Finally, the functional images were acquired with the spin echo imaging sequence epfid2d1, with 2mm isotropic voxel size, TR=800ms, TE=37ms, flip angle=52°.

### Retinotopic mapping stimuli and analysis

A drifting bar stimulus was presented on an MRI compatible LCD screen that was positioned at the end of the scanner bore and viewed by the subject via an angled mirror. The stimulus was composed of black and white checks that contrast reversed at 5 Hz, presented against a mid-gray background. The stimulus was presented within a circular aperture of 21° of visual angle. The bar drifted along the cardinal and oblique orientations, and the sequence of bar positions was played in reverse for the second half of each acquisition. Subjects were asked to monitor a central, black fixation dot during the measurements, and make a bilateral button press when the dot infrequently and briefly changed color to red. The software used to produce the stimuli is publicly available (https://github.com/gkaguirrelab/tomeSpatialStimuli). Subject fixation and alertness were monitored using an in-bore infrared camera (Cambridge Research Systems LiveTrack). Following a pre-registered protocol (https://osf.io/ervrh/), scanning was stopped, and the acquisition repeated, when the subject was observed to lose fixation or become sleepy.

HCP minimal processing pipeline was applied to the functional MRI data (Glasser et al., 2013). The pipeline included corrections for gradient nonlinearity, motion, and phase encoding related distortion at the volumetric level and additional smoothing and vertex area corrections at the surface level. The HCP version of the FSL FIX protocol was then used to further correct the noise in the data (Salimi-Khorshidi et al.,2014; Griffanti et al., 2014).

Population receptive field (pRF) maps were produced from the retinotopic mapping data from each subject (Dumolin & Wandell, 2008). Non-linear model fitting was performed using custom software (https://github.com/gkaguirrelab/forwardModel), derived in part from the analyzePRF toolbox (Kay et al., 2013). The model included a parameter for the timing of the peak of the hemodynamic response function, as the forward and reverse sequence of the stimuli constrains this value. The resulting pRF measurements were then combined with the cortical surface topology derived from the T1 acquisition for each subject within a Bayesian fitting framework (Benson & Winawer, 2018) to produce a final, continuous retinotopic map. From this, we identified the borders of the entire cortical area V1 for each subject.

### Anatomical MRI processing

An unbiased anatomical template was created with the T1w images from all subjects using the multivariate template construction algorithm available in Advanced Normalization Tools (ANTs, Avants et al., 2011). Freesurfer cortical reconstruction and automatic segmentation pipeline was used to define the optic chiasm (Dale et al., 1999), and the thalamic segmentation algorithm was used to define the lateral geniculate nuclei (LGN) on this template (Iglesias et al., 2018). The resulting segmentation masks were used for tractography in a subsequent step. The segmentation was also run on the structural images to find the volume of the LGN for each subject.

### Diffusion MRI Pre-Processing and Fixel-Based Analysis

HCP minimal processing pipeline was also applied to the diffusion images. This pipeline included topup, eddy, and motion corrections (Glasser et al., 2013). For the fixel-based analysis, we opted to use only the highest b-value as fiber density measurements have been shown to be more accurate when b-values at and above 3000 were used for FOD construction (Genc et al., 2020a). Therefore, we extracted b=0 and b=3000 volumes from the preprocessed diffusion images of each subject and continued the rest of the analysis with this reconstructed, single-shell dataset. The images were resampled to 1.25 mm isotropic resolution to increase the anatomical contrast (Dyrby et al., 2014) and white matter (WM), gray matter (GM), and cerebrospinal fluid (CSF) diffusion response functions were calculated for each subject with an unsupervised method (Dhollander et al., 2019). These response functions were averaged across subjects to create population-based response functions for each tissue type. Based on these three tissues, a single-shell, 3-tissue constrained spherical deconvolution method was utilized to calculate FOD maps (Dhollander & Connelly, 2016), using the MRtrix3Tissue package (https://3Tissue.github.io) which is an extension of MRtrix3 (Tournier et al., 2019). The intensity of each FOD map was normalized and an unbiased FOD template was constructed. Fixel-based analysis was performed on this FOD template to obtain whole-brain fiber density (FD), fiber cross section (FC), and the combined fiber density & cross section metric (FDC).

The LGN and optic chiasm masks (previously defined on the T1w template) were linearly registered to the FOD template space. Using these registered masks, optic tract streamlines were calculated for each hemisphere separately with probabilistic parallel transport tractography (Aydogan & Shi, 2021). Prior to this calculation, the optic chiasm mask was dilated to ensure that it covered the entire region adequately, as the initial segmentation tended to underestimate this anatomical structure. Finally, the streamlines were used to crop the whole-brain FD and FC maps and obtained optic tract fixel values. The fixels within the tracts were averaged to obtain a single FD and FC value for each subject.

Due to the large individual variability in the shape and size of human V1, a different method was employed to define optic radiations. Instead of defining them on the FOD template, these were defined on the individual FOD maps. Fixel-based analysis is performed within a template space to allow morphometric measurements. Therefore, following the tractography, the optic radiation streamlines for each subject were converted to volumetric masks and warped to the template FOD space using the warp fields calculated previously as a part of the whole brain fixel analysis. Once the masks were all in the template space, their intersection was calculated, and this intersection mask was used to crop the fixel masks to ensure the final segmentation only included the regions which are present in all 42 subjects.

### Analysis of biometric variation

We collected from each subject their self-reported height and weight. From their T1 image a measure of their intra-cranial volume (ICV) was derived with CAT12 toolbox (http://www.neuro.uni-jena.de/cat/) for SPM (http://www.fil.ion.ucl.ac.uk/spm). As we expected there to be correlated variation in these measures across subjects, these three variables were examined using a principal components analysis. As the scales and the ranges of the input variables were different, the analysis was performed on the correlation matrix rather than the covariance matrix.

The code used for this, and all other analyses reported here is publicly available (https://github.com/gkaguirrelab/retinaTOMEAnalysis).

## Results

We obtained a measure of the volume of the ganglion cell layer of the retina from each of 42 subjects using optical coherence tomography (Figure 1). As described in prior work (Chen et al., 2020) this raw measurement was adjusted for differences in the axial length of the eye, yielding an estimate of individual variation in the number of retinal ganglion cells. Here we will refer to this adjusted measure as “RGC volume”. From these same subjects we obtained a measure of the fiber cross-section (FC) and fiber density (FD) from the optic tracts (Figure 2) and optic radiations (Figure 3). Our primary goal was to examine the relation between individual differences in the retinal measure with individual differences in the fixel properties of post-retinal white matter pathways. First, however, we considered the possibility that variation in overall body size might influence these measures.

**Figure 1.**
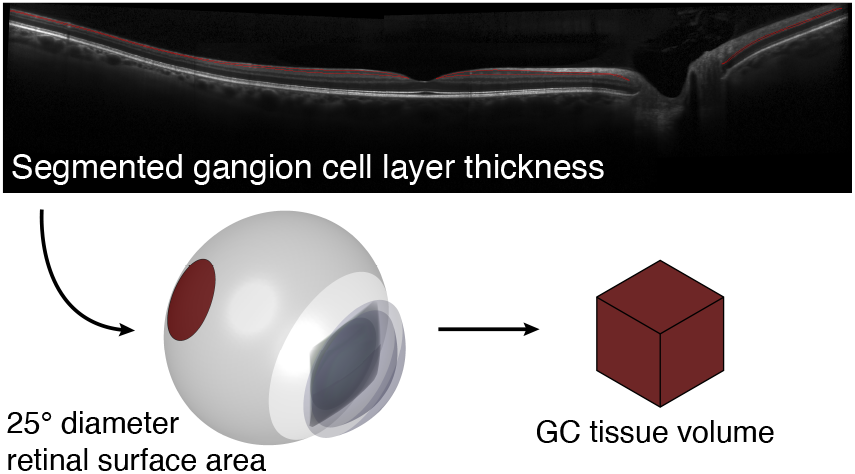
Derivation of RGC volume. In a prior study we obtained OCT images through the fovea along the horizontal and vertical meridians out to 25° radial eccentricity from each of 42 subjects. These images were of sufficient quality to allow segmentation of the ganglion cell layer (red lines) from the inner plexiform layer. The axial length of the eye was also measured, and a model of eye biometry was then used to estimate the total volume of retinal ganglion cell tissue.

**Figure 2.**
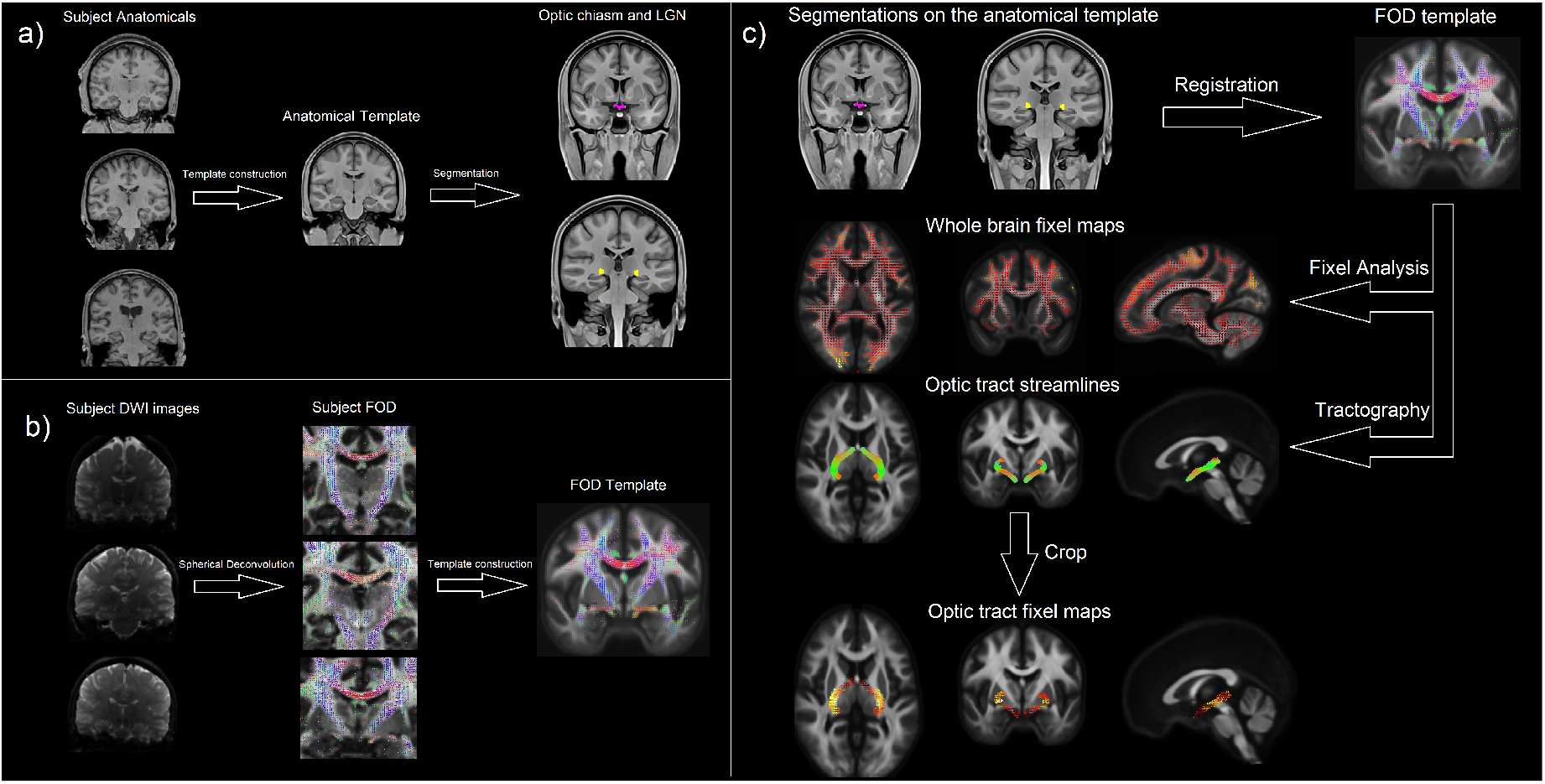
Template construction and optic tract fixel analysis. a) An unbiased anatomical template was derived from the MPRAGE images of all subjects using the ANTs multivariate template construction algorithm. This template was segmented with Freesurfer tools to construct optic chiasm and LGN masks. b) Another unbiased template was built from the FOD images of the same subjects which were calculated with the constrained spherical deconvolution method. c) The masks defined on the anatomical template were registered to the FOD template coordinates and probabilistic tractography between optic chiasm and LGN was performed. Separately, fixel analysis was performed with this FOD template resulting in the whole brain FC and FD fixel maps for each subject. These fixel maps were then cropped with the optic tract streamlines to obtain fixel values from the optic tract.

**Figure 3.**
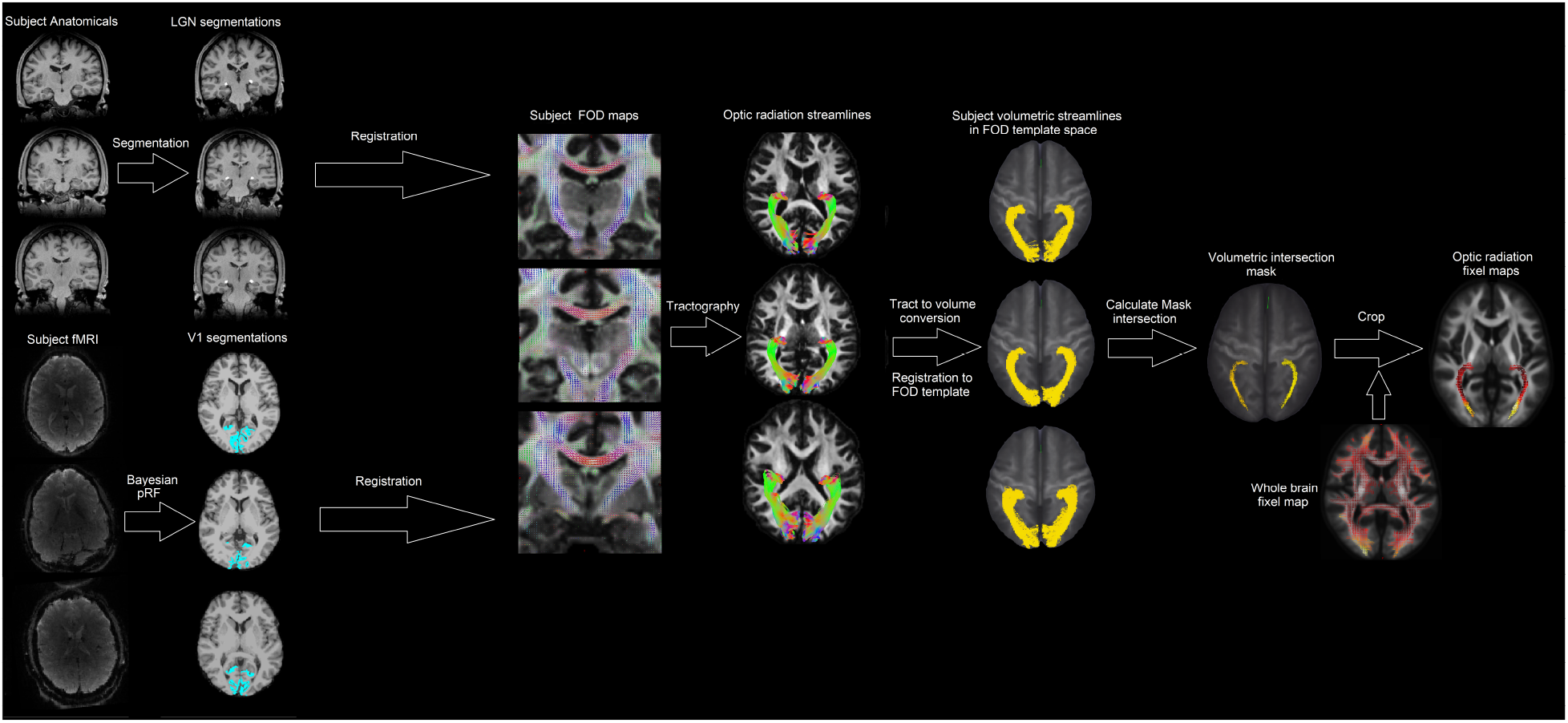
Optic radiation fixel analysis. To calculate the optic radiation streamlines LGN and V1 masks were created using segmentation and pRF analysis for each subject. The segmentations were registered to their respective FOD maps and optic radiation tracts were defined for each subject separately. These tracks were then converted to volumetric masks and registered to the FOD template coordinates using the same warp fields calculated for the fixel analysis. Within the template space, an intersection mask was calculated from all optic radiation streamlines which was then used to extract fixel values from the whole brain fixel maps.

### Individual differences in body size confound visual pathway measures

We obtained measures of height, weight, and intra-cranial volume (ICV) from each of our participants. We used a principal component analysis to summarize these measures and found that the first component accounts for 70% of the variance. We then examined how well this first, “body size” component explains variation in our retinal and optic tract measures. While body size was weakly related to RGC volume (r = 0.28 [95% CI = −0.06, 0.53]), there was a substantial positive correlation with optic tract FC (r = 0.49 [95% CI = 0.21,0.68]), and a negative correlation with optic tract FD (r = −0.56 [95% CI = −0.70, −0.26]) (Figures 4a, b).

**Figure 4.**
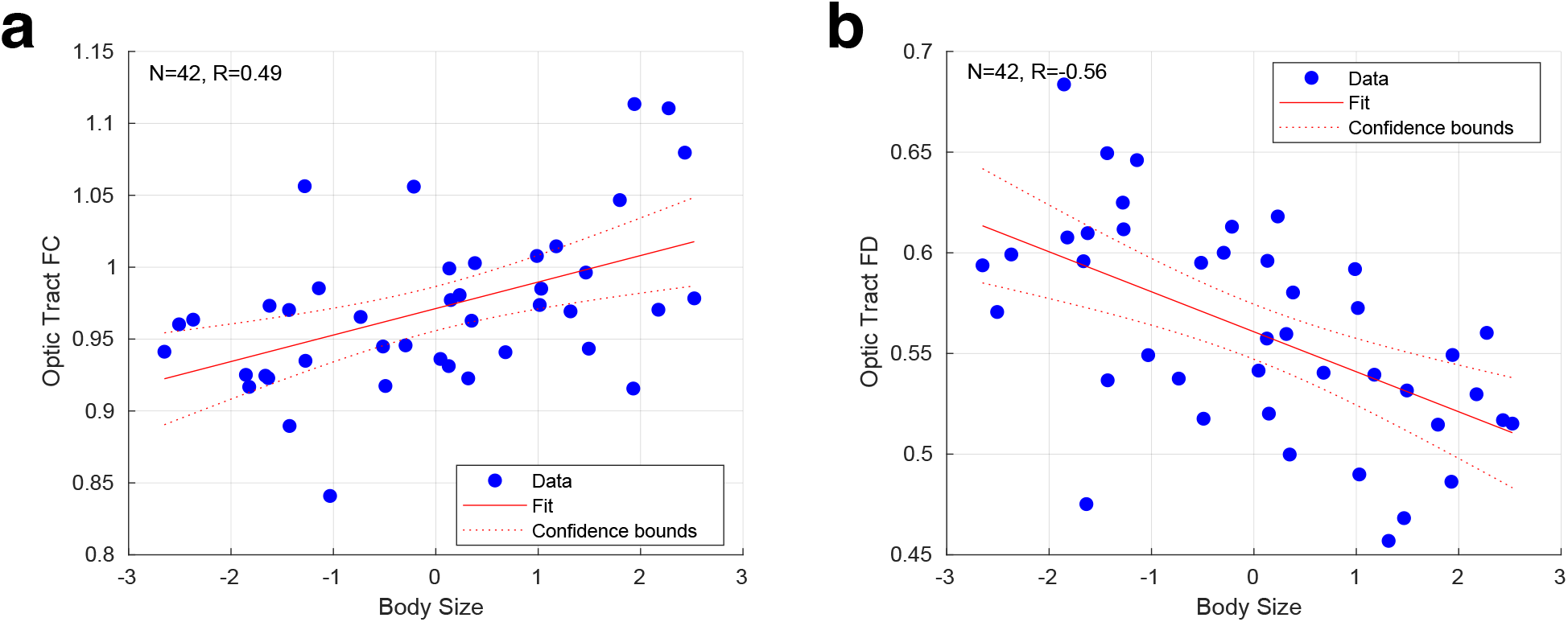
Relationship of size with optic tract fixel measurements. Correlation between the first (“body size”) principal component derived from the biometric measures and a) optic tract FC and b) optic tract FD. The 95% confidence interval of the regression slope is shown.

Our goal is to identify how variation in RGC endowment influences the properties of post-retinal structures. Variation in body size may cause both RGC and anatomical measures to co-vary in a non-selective manner. Therefore, in subsequent analyses, we performed partial regression to remove the linear effect of body size variation prior to examining the relationship between retinal and post-retinal measures.

### RGC volume is correlated with optic tract FC but not FD

After adjusting for body size, we found a significant positive correlation between RGC volume and optic tract FC (Figure 5a; r = 0.37 [95% CI = 0.10, 0.62]), but did not observe this for optic tract FD (Figure 5b; r = 0.06 [95% CI = −0.24, 0.36]). At face value, this result suggests that individual differences in retinal ganglion cell endowment are reflected in the relative size of the optic tract, but not in the diameter or packing density of these axons (after correcting for eye and body size).

**Figure 5.**
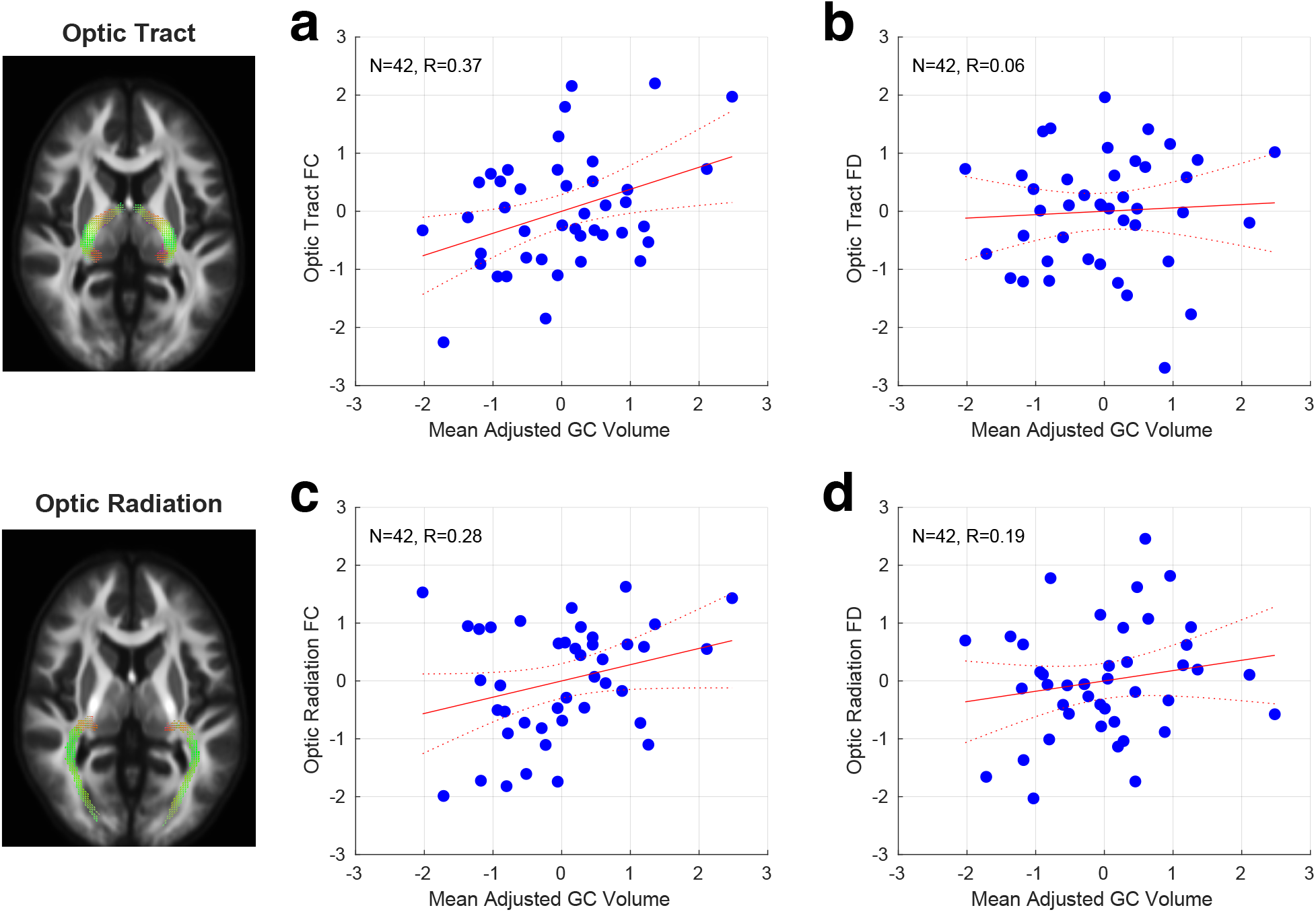
Relationship of RGC volume with fixel measures of the optic tract and radiations. Correlation across subjects between RGC volume and fixel measures. All measures were adjusted for individual differences in overall body size. a) Optic tract FC; b) Optic tract FD; c) Optic radiation FC; d) Optic radiation FD.

Our measure of RGC volume is adjusted, based upon theoretical considerations, for a confounding effect of the axial length of the eye, resulting in a measure that putatively better reflects individual differences in the number of retinal ganglion cells (Chen et al, 2020). While there is reason to believe that variation in axial length is not systematically related to individual differences in the number of retinal ganglion cells, it remains possible that larger eyes may contain larger ganglion cell bodies and associated axons. If so, the raw measure of RGC volume, unadjusted for axial length, may demonstrate a positive relationship with optic tract FD, while the adjusted value does not. To test this idea, we examined the correlation between RGC volume, not adjusted for axial length, and the optic tract measures (while still correcting for overall body size). This unadjusted measure of RGC volume was still not related to optic tract FD (r = 0.05 [95% CI = −0.31, 0.36]). Further, we find that the unadjusted RGC measure has a reduced relationship with optic tract FC (r = 0.21 [95% CI = −0.12, 0.47]).

Next, we considered the possibility that noise in our measurements was sufficient to obscure a true relationship between RGC volume and optic tract FD. To examine this, we obtained the separate value for each measurement type from the left and right side (eye, or hemisphere, as appropriate). Table 1 presents the correlation between left and right measures across subjects. Using these values, and assuming that between-subject differences are substantially larger than within-subject laterality differences, we may calculate a noise-limited ceiling correlation value that would be expected in the presence of an underlying, perfect correlation between different measures. These values were 0.87 and 0.86, respectively, for the relationship between RGC volume and optic tract FC and FD, suggesting that we could have observed a relationship between the retinal measure and optic tract FD if it indeed existed.

**Table 1.** Correlation within measure between sides. A separate value for each measure was obtained from the left and right eye or hemisphere, as appropriate. The correlation across subjects between sides is given, along with the 95% confidence interval obtained by bootstrap resampling.

|  | Left – right correlation |
| --- | --- |
| *adjusted RGC volume* | 0.64 [0.46, 0.79] |
| *Optic Tract FC* | 0.93 [0.88, 0.96] |
| *Optic Tract FD* | 0.91 [0.85, 0.95] |
| *Optic Tract FDC* | 0.92 [0.85, 0.96] |
| *Optic Radiation FC* | 0.86 [0.77, 0.92] |
| *Optic Radiation FD* | 0.76 [0.60, 0.87] |
| *Optic Radiation FDC* | 0.81 [0.61, 0.89] |
| *LGN* | 0.44 [0.08, 0.69] |
| *V1 surface* | 0.81 [0.71, 0.88] |

Our primary analysis made use of the highest b-value diffusion shell, which is thought to increase the relative sensitivity of the FD measure to intra-cellular axonal signals (Genc et al., 2020a). We nonetheless considered the possibility that a different approach to the analysis of the diffusion data would reveal a true relationship between RGC volume and FD. Therefore, we repeated the analysis using all b-values, and a multi-shell, multi tissue, constrained spherical deconvolution (Jeurissen et al., 2014) and re-calculated the fixel metrics. Again, there was no systematic relationship between RGC volume and FD derived from this multi-shell analysis (r = 0.01 [95% CI = −0.29, 0.31).

### RGC volume is not significantly correlated with optic radiation FC or FD

Having found a relationship between RGC volume and optic tract FC, we next asked if this relationship extends to the optic radiations at the next stage of the visual pathway. We identified the LGN in each subject using a template-based segmentation technique, and area V1 using retinotopic mapping, and then measured the FC and FD metrics of the optic radiations using a fixel-based analysis. Neither of these fixel measures were significantly correlated with RGC volume (Figure 5c, d): FC (Figure 5c; r = 0.28 [95% CI = −0.09, 0.57]); FD (Figure 5d; r = 0.19 [95% CI = −0.10, 0.46]).

We considered the possibility that a direct measurement of optic radiation volume in each subject would show a stronger relationship with RGC volume than did the FC measure, which was derived within the across-subject intersection of the optic radiations within the template space. Volume derived from the entire optic radiation streamlines in each subject was also not correlated with RGC volume (r = −0.12 [95% CI = −0.43, 0.20]).

### The sizes of adjacent structures in the visual pathway are well correlated

In addition to the fixel-based measurements of white matter structures, our analysis yielded the volume of the LGN itself, as well as the surface area of V1. We examined the cross-correlation matrix across subjects for the entire set of anatomical measurements. Table 2 (top) provides these correlations for anatomical structures, including the FC measurement of the optic tract and optic radiations. Generally, there are robust correlations between adjacent structures on the visual pathway. These correlations are weaker when measured between more distant structures. This property applies as well to the measurement of RGC volume, which appears to have a decreasing influence upon the visual pathway with distance from the retina.

**Table 2.** Full cross-correlation matrix between structures. The correlation across subjects between each pairing of structures is given. The three separate groupings provide the correlations when white matter properties are measured using FC, FD, or FDC. Values associated with 95% confidence intervals that do not include zero are marked with an asterisk. The calculated noise ceiling correlation value is given in square brackets.

|  | RGC Volume | Optic Tract FC | LGN | Optic Radiation FC |
| --- | --- | --- | --- | --- |
| *Optic Tract FC* | 0.37* [0.87] |  |  |  |
| *LGN* | 0.21 [0.69] | 0.59* [0.77] |  |  |
| *Optic Radiation FC* | 0.28 [0.85] | 0.64* [0.95] | 0.35* [0.75] |  |
| *V1 Surface Area* | 0.06 [0.84] | 0.31* [0.93] | 0.23 [0.74] | 0.53* [0.91] |

|  | RGC Volume | Optic Tract FD | LGN | Optic Radiation FD |
| --- | --- | --- | --- | --- |
| *Optic Tract FD* | 0.06 [0.86] |  |  |  |
| *LGN* | 0.21 [0.69] | 0.05 [0.76] |  |  |
| *Optic Radiation FD* | 0.19 [0.82] | 0.24 [0.91] | -0.21 [0.73] |  |
| *V1 Surface Area* | 0.06 [0.84] | -0.14 [0.92] | 0.23 [0.74] | -0.16 [0.88] |

**Table 2. Full cross-correlation matrix between structures**
|  | RGC Volume | Optic Tract FDC | LGN | Optic Radiation FDC |
| --- | --- | --- | --- | --- |
| *Optic Tract FDC* | 0.22 [0.87] |  |  |  |
| *LGN* | 0.21 [0.69] | 0.31* [0.77] |  |  |
| *Optic Radiation FDC* | 0.32 [0.83] | 0.42* [0.93] | 0.15 [0.74] |  |
| *V1 Surface Area* | 0.06 [0.84] | 0.05 [0.93] | 0.23 [0.74] | 0.37* [0.89] |

The relationship between RGC volume and V1 surface area is notably low. This analysis was conducted using the absolute surface area of V1 in each subject. We considered that the proportion of the cortical surface devoted to primary visual cortex, as opposed to raw area, might be more strongly related to the retinal measure. However, even when expressing the size of area V1 as a proportion of total surface area, there was no relationship with RGC volume (r = −0.03 [95% CI = −0.31, 0.25]).

We repeated the set of cross-correlation analyses using the FD white matter measure, as well as the fiber density & cross section metric (FDC), which is proposed to be a more comprehensive assessment of total intra-axonal volume (Raffelt et al., 2017). In all cases, the FD-associated measures are less related to adjacent or distant visual pathway anatomy (Table 2, middle and bottom).

### Optic tract, but not optic radiation, FC and FD values are correlated across subjects

Overall, we find that adjusted RGC volume is related to optic tract FC, and that FC values from the optic tract and optic radiations are related to each other and to adjacent structures in the visual pathway. In contrast, we don’t observe any of these relations for FD. This might raise the concern that FD is an unreliable measurement or is unrelated to an underlying biological property of the tissue. Mitigating these concerns, we find that FD is strongly correlated between left and right hemisphere structures across subjects, arguing against the measurement being subject to high within-session noise. After adjusting for body size, we also find that optic tract FC is related to optic tract FD across subjects (r = 0.40 [95% CI = 0.22, 0.55]), although this relationship is not present for the optic radiations (r = 0.03 [95% CI= −0.28, 0.37]).

## Discussion

We find that individual variation in the overall volume of the retinal ganglion cell layer of the eye has limited impact upon the structural properties of the post-retinal visual pathway. RGC variation influences the caliber of the optic tracts, with decreasing and non-significant correlations with subsequent visual pathway structures. We are able to interpret our findings of minimal or null effects as meaningful, given the good reproducibility of our measurements and the observation of many other strongly related elements of anatomical variation.

### Fixel measures in the optic tract

A goal of our work was to evaluate if the fixel analysis approach is sensitive to biological variation in a healthy population. Each retinal ganglion cell gives rise to a single axon within the optic tract. Fixel analysis yields estimates of the cross-sectional area of a fiber bundle (FC), and of the proportion of intra-axonal volume (FD). Nominally, fixel measures of the optic tract should reflect individual variation in the number of retinal ganglion cells in the eyes, either in FC, FD, or their combination. We find that, while individual retinal variation is correlated with FC, it has effectively no influence upon measured FD. This remains true with or without adjustment for variation in the overall size of the eye. Within the optic tract itself, however, we find that individual variation in FC and FD are positively correlated.

We interpret this set of findings as showing that people with a greater endowment of retinal ganglion cells have a commensurately larger optic tract, but that this increase in the caliber of the fiber bundle results in no net change in the density of axonal packing. An independent, individual difference varies axonal diameter, leading to yoked changes in optic tract FC and FD.

Our analyses were conducted after correcting for the confounding effects of variation in overall body size. We found a strong positive correlation between body size and optic tract FC. This relationship is sensible, and a similar effect of variation in intracranial volume upon central nervous system FC measurements has been reported (Pannek et al., 2018; Smith et al., 2019). Increased body size in mammals is associated with both an increase in axon diameter, and a decrease in axon packing density within nerves (Wang et al., 2008). We found a strong negative correlation between body size and FD, suggesting that a decrease in packing density is the predominant effect of body size variation in the optic nerve. Similar, negative relationships of body size and FD have been previously reported for other brain locations (Choy et al., 2020).

Similar to many others studies, our fixel measurements were derived from diffusion MR data collected with a gradient b-value of 3000. There is evidence that even higher b-values can enhance the contribution of intra-axonal volume to the FD measure (Genc et al., 2020a). It is therefore possible that a subtle relationship between FD and other anatomical structures, missed in this study, might be revealed using different imaging parameters.

### RGC influence declines along the visual pathway

We find that the influence of RGC volume on the visual pathway declines steadily as we move past the optic tract. At the level of the primary visual cortex, there is no relationship between V1 surface area and individual variation in our retinal ganglion cell measure. We do find, however, strong positive relationships in the size of adjacent elements of the anatomical pathway, similar to prior work (Kupfer et al., 1967; Andrews et al., 1997). The current and prior results suggest that there are multiple, local influences upon variation in the size of visual pathway structures. For example, the relative size of the visual cortex has a heritable component (Bakken et al., 2012; Yoon et al., 2019), although there is limited study of genetic influences elsewhere along visual pathway.

In a recent study, Miyata et al., 2022 investigated how V1 surface area is related to measures of visual pathway white matter using diffusion tensor (DTI; Basser et al., 1994) and neurite density and orientation dispersion imaging (NODDI; Zhang et al., 2012). They reported a small but significant negative correlation between the fractional anisotropy (FA) of the optic tracts and the surface area of the V1. In the supplementary materials we present a DTI analysis of our data. Similar to Miyata and colleagues, we find that optic tract FA (and MD) has a modest negative correlation with V1 surface area. However, this correlation is not present after accounting for variation in body size confound, raising the possibility that a similar effect may be present in the data examined by Miyata and colleagues.

We studied here a measure of individual variation in RGC tissue, integrated over eccentricity and angular position, and did not observe a relationship with variation in the area of V1 cortex. It is possible that individual variation in the distribution (as opposed to overall quantity) of retinal cell populations over eccentricity or polar angle may be reflected in the visual cortex. Indeed, recent work has shown that polar angle variation in group and individual measures of perception are related to variation in cortical magnification across area V1 (Benson et al., 2021; Himmelberg, et al., 2021; Kupers et al., 2022).

### Relation to prior work in clinical populations

We are unaware of prior efforts to relate individual variation in the healthy retina to diffusion measures of the visual pathway. Diffusion imaging has been related, however, to group differences between healthy controls and patients with ophthalmologic disease, and to individual variation in measures of retinal injury. Most relevant to the current study, Haykal and colleagues (2019) found a significant decrease in FC and FD measured in the optic tracts (and optic radiations) of patients with glaucoma as compared to healthy controls. In comparison to healthy variation, glaucoma causes a loss of RGCs, resulting in axonal degeneration within an existing fiber bundle. It is sensible that such an injury would cause a reduction of both the overall size and the intra-axonal proportion of the optic tract.

### Conclusions

We find that a fixel-based measurement of the cross-sectional area of the optic tract appropriately reflects individual variation in retinal ganglion cell endowment. While we did not observe this relationship for the FD metric, we suspect that the measure is nonetheless a valid estimate of the proportion of intra-axonal area, given the across-subject relationship we observe between FD and FC within the optic tract, and prior work in patient populations (Haykal 2019). Our work also highlights the importance of considering how overall variation in body size, and specific variation in the properties of a sensory organ (i.e., the eye), can influence measurements of across subject variation of the central nervous system.

## Supporting information

Supplementary analysis

