## Supplementary analysis for "Retinal ganglion cell endowment is correlated with optic tract fiber cross section, not density"

### Supplementary material

We conducted an additional analysis to examine how variation in diffusion tensor imaging (DTI) measures relate to RGC volume. DTI-based fractional anisotropy (FA) and mean diffusivity (MD) metrics were measured for the optic tracts and optic radiations in each subject. A DTI model was fitted to the diffusion images using FSL's "dtifit" function after the images were processed with the HCP minimal preprocessing pipeline. The optic tract and optic radiation masks that were previously defined on the FOD template coordinates for the fixel analysis were warped to the resulting FA and MD maps using the inverse of the registration matrices that had previously been generated to warp each FOD map from each subject to template FOD coordinates. Using these masks, the average FA and MD values were extracted from the optic tract and optic radiations for each subject. The table below provides the correlation of FA and MD with other structures on the visual pathway, with and without correction for variation in body size.

Measures for which the 95% confidence interval did not include zero are marked with an asterisk, and the noise ceiling correlation calculated for the comparison is given in square brackets.

The work of Miyata and colleagues (2022) examined the relationship between optic tract FA, and V1 surface area. This correlation moves from -0.23 to -0.06 after correction for body size. With correction for body size, optic tract FA and MD have non-zero correlations with the corresponding measures from the optic radiations.

### Uncorrected for body size

| | RGC Volume | Optic Tract FA | LGN | Optic Radiation FA |
|---|---|---|---|---|
| *Optic Tract FA* | -0.04 [0.85] | | | |
| *LGN* | 0.21 [0.69] | -0.01 [0.75] | | |
| *Optic Radiation FA* | 0.05 [0.75] | 0.23 [0.82] | -0.15 [0.67] | |
| *V1 Surface Area* | 0.06 [0.84] | -0.23 [0.91] | 0.23 [0.74] | 0.01 [0.81] |

| | RGC Volume | Optic Tract MD | LGN | Optic Radiation MD |
|---|---|---|---|---|
| *Optic Tract MD* | -0.22 [0.82] | | | |
| *LGN* | 0.21 [0.69] | -0.38* [0.73] | | |
| *Optic Radiation MD* | 0.18 [0.85] | 0.43* [0.90] | -0.09 [0.75] | |
| *V1 Surface Area* | 0.06 [0.84] | -0.34* [0.88] | 0.23 [0.74] | 0.09 [0.91] |

### Corrected for body size

| | RGC Volume | Optic Tract FA | LGN | Optic Radiation FA |
|---|---|---|---|---|
| *Optic Tract FA* | 0.07 [0.85] | | | |
| *LGN* | 0.21 [0.69] | 0.12 [0.75] | | |
| *Optic Radiation FA* | 0.03 [0.75] | 0.29* [0.82] | -0.18 [0.67] | |
| *V1 Surface Area* | 0.06 [0.84] | -0.06 [0.91] | 0.23 [0.74] | -0.03 [0.81] |

| | RGC Volume | Optic Tract MD | LGN | Optic Radiation MD |
|---|---|---|---|---|
| *Optic Tract MD* | -0.06 [0.82] | | | |
| *LGN* | 0.21 [0.69] | -0.24 [0.73] | | |
| *Optic Radiation MD* | 0.20 [0.85] | 0.50* [0.90] | -0.08 [0.75] | |
| *V1 Surface Area* | 0.06 [0.84] | -0.07 [0.88] | 0.23 [0.74] | 0.13 [0.91] |
